# Sex-Specific Regulation of Endothelial-to-Mesenchymal Transition (EndMT) During Atherosclerosis

**DOI:** 10.64898/2026.02.07.704593

**Authors:** Kelsey Watts, Katyayani Sukhavasi, Rebecca Hernandez, Alexia Wallace, Daniek Kapteijn, Ernest Diez Benavente, Michal Morky, Noah Perry, Husain Ahammad Talukdar, Johan L M Björkegren, Karen Reue, Hester den Ruijter, Mete Civelek

## Abstract

Sex differences in atherosclerotic plaque biology underlie clinically distinct manifestations of acute coronary syndromes, yet the molecular mechanisms driving these differences remain incompletely understood. Endothelial-to-mesenchymal transition (EndMT) is increasingly recognized as a contributor to plaque remodeling, but whether EndMT is regulated in a sex-specific and stage-dependent manner across atherosclerosis has not been systematically examined. Here, we integrated bulk RNA sequencing with single-cell transcriptomic analyses in endothelial cells and human atherosclerotic plaques to define sex-specific EndMT regulation across healthy and advanced-stage disease. Using trajectory analyses in endothelial cells undergoing EndMT, we observed pronounced sex-specific regulation in healthy endothelium, with females exhibiting stronger early EndMT activation, whereas endothelial cells derived from atherosclerotic plaques displayed markedly attenuated sex differences with disease progression. Consistent sex-divergent pseudotime trajectories were observed in human carotid plaque endothelial single-cell RNA-seq data, with females showing greater EndMT activation at earlier stages and males at later stages. Together, these findings support a stage-dependent model of sex-specific EndMT regulation, indicating that the functional consequences of EndMT are highly context dependent and may differ across early and late disease stages. Integration of these datasets prioritized high-confidence sex-specific EndMT regulators, including *COL4A1, PECAM1, CD151, JAG1, FN1, NEDD9, PODXL, MAFB, PROCR*, and *CDH13*, providing a mechanistic framework to explain clinically observed sex differences in plaque biology and to guide targeted functional interrogation.

## Introduction

Acute Coronary Syndrome (ACS) occurs when blood flow to the heart is suddenly reduced. The two primary types of ACS are plaque rupture, accounting for 50% of all cases, and plaque erosion, which accounts for 40%^1^. Plaque rupture typically involves a thin fibrous cap overlying a large necrotic core, leading to thrombus formation upon rupture, while plaque erosion is characterized by endothelial loss over an intact, thick fibrous cap. Clinically, plaque rupture is more prevalent in men and underlies the majority of myocardial infarctions, whereas plaque erosion is more common in women, particularly younger women^2^. These distinct pathologies reflect underlying sex differences in plaque biology, as females generally exhibit more stable, fibrotic plaques with thicker caps, whereas males tend to develop lipid-rich plaques with larger necrotic cores^3^. Although these sex-specific features of plaque biology are well established, the mechanisms driving such divergent phenotypes remain poorly understood, and few studies have directly examined the molecular pathways contributing to these differences in atherosclerotic disease progression.

Emerging evidence suggests that endothelial-to-mesenchymal transition (EndMT) may contribute to these sex differences in plaque biology^4–7^. EndMT describes a process by which endothelial cells lose their characteristic markers and functions and acquire mesenchymal properties, including increased migratory and contractile capacity. This phenotypic plasticity is now recognized as a significant driver of vascular remodeling and inflammation in atherosclerosis^8,9^. In murine lineage tracing models, EndMT-derived cells have been identified within atherosclerotic lesions, contributing to both destabilizing processes such as neovascularization and protective remodeling that thickens the fibrous cap^10–13^. There is emerging evidence suggesting that approximately one quarter of the cells forming the fibrous cap are of endothelial origin rather than smooth muscle lineage, suggesting a significant role for endothelial plasticity in determining plaque phenotype^14^.

There is no standardized approach for modeling EndMT in vitro, however, biochemical induction using transforming growth factor beta (TGFβ), particularly the TGFβ2 isoform, combined with an inflammatory cytokine such as Interleukin-1β (IL-1β) or Tumor Necrosis Factor α (TNFα), is among the most commonly employed strategies^9,15,16^. TGFβ promotes EndMT primarily through activation of canonical SMAD2/3 signaling, which drives transcriptional programs that repress endothelial identity and induce mesenchymal gene expression^11^. In parallel, TGFβ also engages non-canonical pathways, including MAPK, PI3K/AKT, Rho-like GTPase, and Notch signaling that contribute to cytoskeletal remodeling, increased motility, and extracellular matrix deposition, collectively reinforcing the endothelial-to-mesenchymal transition^8^. TNFα acts synergistically by activating NFκB-dependent signaling, which promotes inflammatory gene expression and further amplifies the transition^17,18^. Because TNFα/NFκB activity exhibits robust sex-specific regulation, examining the combined influence of TGFβ and TNFα may reveal mechanisms driving differential EndMT activation between males and females^19,20^.

Assessment of EndMT relies on a combination of complementary molecular, morphological, and functional readouts rather than a single definitive marker. EndMT is commonly evaluated through coordinated changes in gene and protein expression, including downregulation of endothelial markers (e.g., *PECAM1, CDH5, VWF*) and induction of mesenchymal-associated genes (e.g., *FN1, ACTA2, TAGLN*), alongside morphological transitions from a cobblestone endothelial phenotype to an elongated, spindle-like morphology^8^. In some experimental contexts, functional assays such as migration, contractility, extracellular matrix deposition, or barrier integrity are also used to capture EndMT-associated phenotypic changes^21^. Consistent with prior studies, we assess EndMT using integrated transcriptional profiling across time, immunofluorescence-based marker expression, and morphological changes, enabling evaluation of partial and stage dependent EndMT programs.

We hypothesized that females exhibit greater activation of EndMT pathways, contributing to enhanced fibrous cap formation and overall plaque stability. To investigate potential sex-specific regulation of EndMT, we induced EndMT in human endothelial cells derived from two vascular sources, healthy aortic endothelium and atherosclerotic carotid lesions, and performed bulk RNA-sequencing over multiple time points following TGFβ and TNFα stimulation to compare transcriptional responses between sexes. To validate these findings in vivo, we analyzed human single-cell RNA-sequencing datasets of atherosclerotic carotid plaques (Figure 1A). This integrated approach enabled identification of sex-specific EndMT signatures and elucidated how differential pathway activation over time might contribute to the distinct structural and clinical features of atherosclerotic plaques observed between men and women.

**Figure 1:**
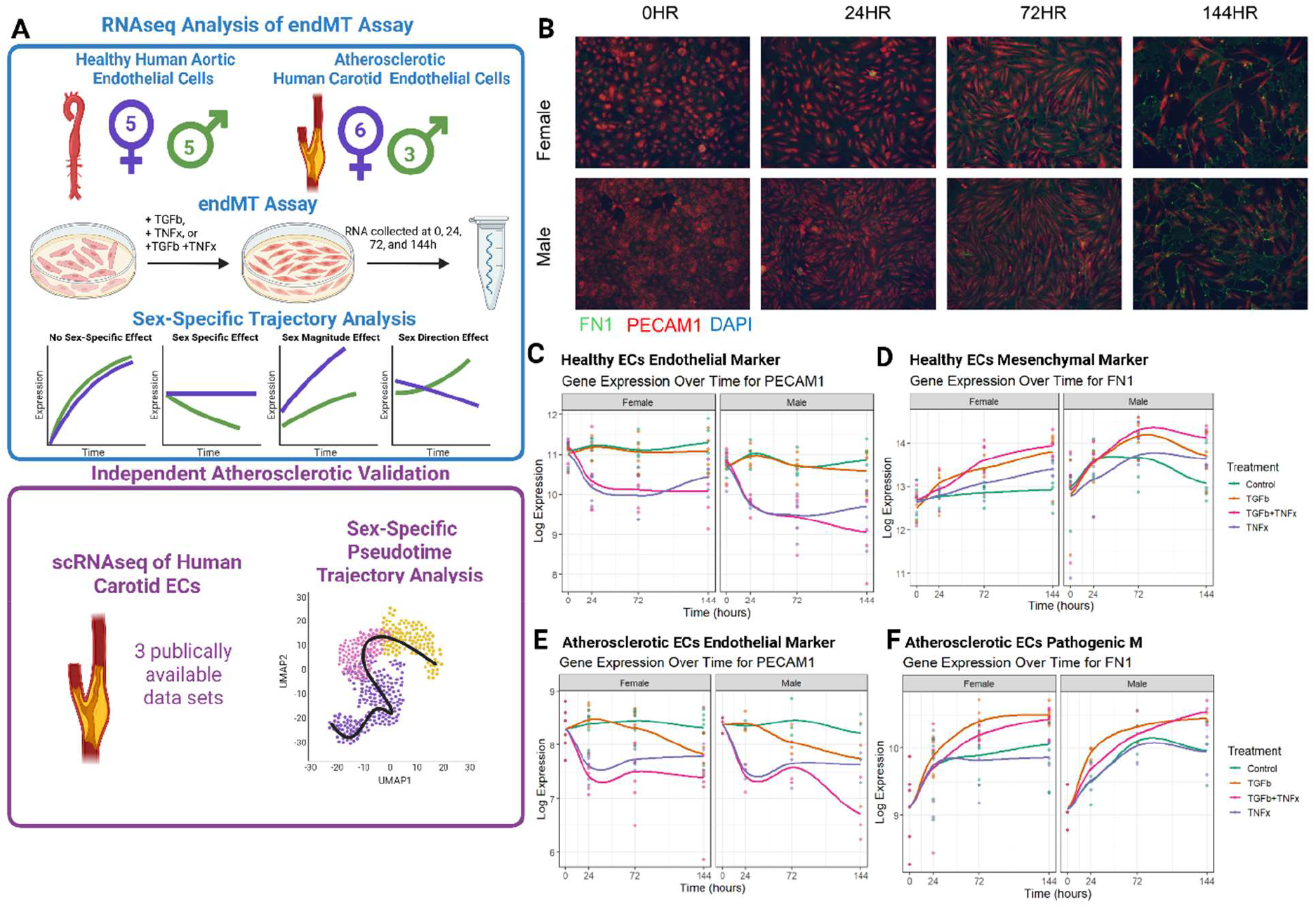
Experimental framework for defining sex-specific regulation of endothelial-to-mesenchymal transition (EndMT). (A) Overview of the integrated workflow. Sex-specific trajectory analysis begins with bulk RNA-seq from an in vitro EndMT assay performed on male and female human endothelial cells (ECs) derived from healthy aortic and atherosclerotic carotid vascular beds. (B) Representative immunofluorescence images from the in vitro EndMT assay showing PECAM1 and FN1 as endothelial and mesenchymal markers, respectively. (C– D) *PECAM1* (C) and *FN1* (D) expression over time in healthy ECs during the EndMT assay. (E– F) *PECAM1* (E) and *FN1* (F) expression over time in atherosclerotic ECs during the EndMT assay.

## Methods

### Endothelial Cell Isolation and Characterization

Primary endothelial cells (ECs) were isolated from the ascending aorta of healthy heart transplant donors (5 male, 5 female), as previously described^22^. Donor metadata (e.g., age, comorbidities, or medications) were not available for these lines; however, given that the average age of heart donation typically ranges from 22–41 years, female donors were likely premenopausal^23^. Donor sex was confirmed by expression of *XIST* (Supplemental Figure S1A).

Plaque ECs (3 males, 6 females) were isolated from fresh atherosclerotic plaque tissues obtained from patients undergoing carotid endarterectomies from the Athero-Express Biobank as previously described^24^. Metadata for these patients can be found in Supplemental Table 1.

**Table 1:** Prioritization of sex-specific EndMT regulators supported across orthogonal human and mouse datasets. Filled circles indicate support for each gene across human and mouse transcriptomic datasets, genetic associations, biological annotations, and network modules. Weighted scores reflect cumulative support across transcriptomic, genetic, and network-level evidence for the top 10 sex-specifc EndMT regulators.

| Top Prioritized Genes Supported Across Orthogonal Datasets |  |  |  |  |  |  |  |  |  |  |  |  |
| --- | --- | --- | --- | --- | --- | --- | --- | --- | --- | --- | --- | --- |
|  | Human Transcriptomics |  |  | Mouse Transcriptomics |  | Genetics |  | Biology |  | Network |  | Weighted Score |
|  | Healthy RNA-seq (0.75) | Pathogenic RNA-seq (0.75) | Human scRNA-seq (1.0) | FCG Gonadal (0.5) | FCG Chromosomal (0.5) | Sex-biased CAD GWAS (1.25) | CAD Association (1.0) | EC Specificity (0.25) | EndMT Pathway (0.25) | Yellow Module (0.25) | Purple Module (0.25) |  |
| COL4A1 |  |  | ● |  | ● | ● | ● |  | ● |  | ● | 4.25 |
| PECAM1 |  |  | ● |  | ● |  | ● | ● | ● | ● |  | 4.00 |
| CD151 | ● |  | ● |  |  |  | ● | ● |  |  |  | 3.50 |
| JAG1 | ● | ● |  | ● |  |  | ● |  | ● |  | ● | 3.50 |
| FN1 |  |  | ● | ● |  |  | ● |  | ● |  |  | 3.25 |
| NEDD9 | ● |  | ● |  |  |  | ● |  |  |  | ● | 3.25 |
| PODXL |  |  | ● | ● | ● |  | ● |  |  |  |  | 3.25 |
| MAFB | ● |  |  |  | ● |  | ● | ● |  |  |  | 3.00 |
| PROCR | ● |  | ● | ● | ● |  | ● |  |  |  |  | 3.00 |
| CDH13 |  |  | ● |  | ● |  | ● |  |  |  |  | 3.00 |

### EndMT in vitro Assay

All endothelial cell lines underwent an EndMT assay to induce mesenchymal transition. This assay was optimized for primary ECs through previously described experiments^7,24,25^. All cells used were between 3-10 passages. Cells were seeded on gelatin-coated (0.2% porcine gelatin, Sigma Aldrich) 12-well plates and maintained at 37°C with 5% CO_2_ until reaching at least 80% confluency. At that point, regular media was replaced with phenol-free EC medium MV (PromoCell) supplemented with 5% charcoal-stripped FBS (Neuromics), and cells were serum/hormone-starved for 24 hours. On day 0, RNA was collected using Qiagen RNeasy kits, and media was refreshed with either control conditions (phenol-free EC medium MV + 5% charcoal-stripped FBS) or stimulation conditions consisting of 25 ng/mL TNFα (Miltenyi Biotec), 10 ng/mL TGF-β2 (PeproTech), or the combination of 25 ng/mL TNFα + 10 ng/mL TGF-β2. RNA was subsequently collected at 24, 72, and 144 hours. Treatment media was refreshed every 48 hours, and cell viability and morphology were monitored throughout using light microscopy.

### Immunofluorescence Staining of EndMT Assay

EndMT-treated endothelial cells were fixed in 4% paraformaldehyde at 0, 24, 72, and 144 hours for morphological assessment by immunofluorescence. Cells were then permeabilized with 0.1% Triton X-100 in PBS, and blocked with 10% goat serum (Sigma-Aldrich). Immunostaining was performed using primary antibodies against PECAM1 (rabbit; Invitrogen MA5-32126; 1:300) and FN1 (mouse; Invitrogen MA5-11981; 1:500), followed by species-appropriate Alexa Fluor–conjugated secondary antibodies (Alexa Fluor 594 goat anti-rabbit; Invitrogen A11001 and Alexa Fluor 488 goat anti-mouse; Invitrogen A11012; 1:1000). Nuclei were counterstained with DAPI (Invitrogen; 1:10,000). Images were acquired using an EVOS M7000 microscope at 20× magnification with identical exposure and gain settings across all conditions.

### RNA Sequencing

Following collection during the EndMT assay, all healthy donor RNA was stored at −80°C until sequencing library preparation. Each donor-derived EC line was treated as an independent biological replicate; no technical replicates were pooled for RNA-seq analyses. RNA from healthy ECs was processed by Admera using the NEBNext Ultra II Directional RNA Library Prep Kit with poly(A) selection. Libraries were sequenced on an Illumina platform using 2 × 150 bp paired-end reads, generating approximately 50 million paired-end reads per sample (25 million reads per direction). Raw paired-end FASTQ files were trimmed using Trimmomatic v0.39 to remove adapters and low-quality bases (ILLUMINACLIP: TruSeq3-PE, SLIDINGWINDOW:4:20, LEADING:3, TRAILING:3, MINLEN:25). Read quality was assessed with FastQC v0.11.5 on all trimmed files. Trimmed reads were aligned to the human GRCh38 reference genome using STAR v2.7.11b with GENCODE v38 annotations, generating coordinate-sorted BAM files and transcriptome-aligned BAM files. Alignment statistics were computed using samtools flagstat. Gene-level expression values were quantified using RSEM in paired-end mode with the STAR transcriptome-aligned BAM files and an RSEM reference built from GRCh38/GENCODE v38. RSEM produced expected counts and TPM values for downstream differential expression analysis.

Atherosclerotic RNA library preparation was performed, adapting the CEL-Seq2 protocol for library preparation. The initial reverse-transcription reaction primer was designed as follows: an anchored polyT, a unique 6bp barcode, a unique molecular identifier (UMI) of 6bp, the 5’ Illumina adapter and a T7 promoter. Complementary DNA was used for in vitro transcription reaction (AM1334; Thermo-Fisher). The resulting amplified RNA (aRNA) was fragmented and cleaned. RNA yield and quality were checked by Bioanalyzer (Agilent).

cDNA library construction was initiated according to the manufacturer’s protocol, with the addition of randomhexRT primer as a random primer. PCR amplification was performed with Phusion High-Fidelity PCR Master Mix with HF buffer (NEB, MA, USA) and a unique indexed RNA PCR primer (Illumina) per reaction. Library cDNA yield was checked by Qubit fluorometric quantification (Thermo-Fisher) and quality by Bioanalyzer (Agilent). Libraries were sequenced on the Illumina NovSeq 6000 platform (Utrecht Sequencing Facility).

Upon sequencing, fastq files were de-barcoded and split for forward and reverse reads. The reads were demultiplexed and aligned to human cDNA reference (GRCh38.p13/ENSEMBL_GENES_108) using the BWA (0.7.13) by calling ‘bwa aln’ with settings -B 6 -q 0 -n 0.00 -k 2 -l 200 -t 6 for R1 and -B 0 -q 0 -n 0.04 -k 2 -l 200 -t 6 for R2, ‘bwa sampe’ with settings -n 100 -N 100. Multiple reads mapping to the same gene with the same unique molecular identifier (UMI, 6bp long) were counted as a single read.

### Sex-Specific Trajectory Analysis

Differential expression analysis was performed in R using edgeR and limma. Expected counts from RSEM were merged with sample metadata, and only genes present in both the healthy and atherosclerotic EC datasets were retained for downstream analysis. We modeled sex and time using a design matrix of the form ∼ 0 + sex * time + donor, treating donor as a blocking factor. Lowly expressed genes were filtered with filterByExpr, and library sizes were normalized using the TMM method. Voom was used to generate voom-transformed log2 counts with precision weights, and duplicateCorrelation was applied to account for within-donor correlation before fitting linear models with lmFit. We then tested sex-specific temporal effects using contrast matrices in limma, with the primary contrast representing an hour-weighted sex difference in response over time (male vs. female AUC-style contrast across 24, 72, and 144 hours relative to 0 hours). P values were moderated with empirical Bayes, and results were reported as log2 fold change, moderated t statistic, and FDR-adjusted p values. Unless otherwise specified, an FDR threshold of 0.25 was used to prioritize candidate genes in discovery-oriented analyses, consistent with exploratory multi-omic integration frameworks. For visualization and pathway analysis, volcano plots were generated from the AUC contrast.

Pathway enrichment analysis was performed in R using fgsea on the limma AUC male vs. female contrast. Genes were ranked by moderated t-statistic, and Hallmark gene sets were obtained via msigdbr (category H). Enrichment was tested using 10,000 permutations, a minimum gene set size of 15, and a maximum of 500 genes. Because the contrast was defined as male minus female, normalized enrichment scores (NES) greater than zero reflect pathways more enriched in males, whereas NES less than zero reflect pathways more enriched in females. Significant pathways were defined as FDR-adjusted p < 0.05, and the top enriched male- and female-associated Hallmark pathways were visualized using bar plots. Because no EndMT-specific gene ontology exists, EMT gene sets are used as a proxy for mesenchymal activation in endothelial cells, consistent with prior literature^8,26^.

### scRNAseq Pseudotime Analysis

Publicly available human carotid endothelial scRNA-seq datasets (Athero-Express^27^, Pan et al.^28^, and STARNET^6^) were integrated in Seurat to generate a unified dataset for sex-specific pseudotime trajectory analysis. UMAP embeddings and SNN clustering were computed using standard Seurat workflows, and clusters were manually annotated based on endothelial, inflammatory, mesenchymal, and smooth muscle-like marker expression. Non-endothelial clusters were removed, and the remaining EC states were converted into a Monocle3. Trajectory inference was performed using EC1 as the root state. Gene expression was modeled using ∼ sex + pseudotime + sex:pseudotime to identify genes associated with pseudotime progression, sex differences, and their interaction (FDR< 0.25). For pathway-level interpretation, pseudotime was divided into early, mid, and late bins; within each bin, all genes were ranked by the signed statistic derived from the sex term and analyzed using Hallmark gene set enrichment (fgsea).

### Weighted Gene Co-expression Network Analysis (WGCNA)

WGCNA was performed on the healthy female EndMT RNA-seq time course. Only samples from EndMT-inducing conditions (TGFβ + TNFα) at 24, 72, and 144 hours were included to focus network construction on transcriptional programs associated with EndMT progression. A signed network was used so that only positively correlated gene pairs contributed to module detection. A soft-thresholding power was selected using pickSoftThreshold, followed by construction of a signed adjacency matrix, topological overlap matrix (TOM), and hierarchical clustering of genes based on TOM dissimilarity. Modules were identified using dynamic tree cutting (minimum module size = 30; deepSplit = 2) and merged at a cut height of 0.25. Module eigengenes (MEs) were defined as the first principal component of each module. EndMT relevance was assessed using the MSigDB Hallmark EMT gene set. Modules were prioritized using an EndMT module score defined as |cor(ME, EMT)| × −log10(EMT FDR). EMT enrichment p-values were adjusted using the Benjamini–Hochberg procedure.

To assess module robustness across sex and disease state, module preservation was evaluated using the healthy female network as the reference and tested in healthy male, atherosclerotic female, and atherosclerotic male EndMT time-course datasets using the WGCNA modulePreservation framework (200 permutations; signed network). Preservation was summarized using Zsummary, with standard thresholds applied (Zsummary < 2, low; 2–10, moderate; ≥10, strong preservation). To account for sex imbalance in the atherosclerotic dataset, a balanced resampling module preservation analysis was performed for selected EndMT-associated modules. Equal numbers of male and female atherosclerotic samples were randomly selected, and module preservation was recalculated using the healthy female network as the reference. This procedure was repeated for 50 resampling iterations, with 20 permutations per iteration. EndMT-associated modules were further evaluated for enrichment of Hallmark EMT genes using one-sided Fisher’s exact tests against the WGCNA gene universe, with Benjamini–Hochberg FDR correction applied across modules; enrichment results with FDR < 0.05 were considered statistically significant.

### Weighted Gene Prioritization

Genes were prioritized to define top sex-specific regulators using a combination of newly generated datasets and relevant published literature. Because a relaxed FDR threshold (0.25) was used during discovery-oriented analyses, genes were not interpreted based on significance in any single dataset but were prioritized only when supported by convergent evidence across independent, orthogonal datasets. Datasets were weighted according to their relevance to human sex-specific cardiovascular biology, with higher weights assigned to evidence derived from human endothelial cells or human genetic association studies. Genes identified as significant in bulk RNA-seq analyses of healthy and atherosclerotic human endothelial cells under the sex-specific trajectory analysis TGFβ+TNFα condition were assigned a moderate weight (0.75). Genes identified from sex-specific pseudotime analysis of human scRNA-seq data were weighted more heavily (1.0), reflecting their in vivo relevance. Magenta module genes identified by WGCNA were included as supportive network-level evidence and assigned a lower weight (0.25).

Additional information from the literature was incorporated to further prioritize this list of candidate genes. Genes identified in sex-biased coronary artery disease (CAD) genome-wide association studies (GWAS) were assigned the highest weight (1.25), while independent CAD associations were weighted strongly at 1.0^29,30^. Additional annotations, including endothelial specificity (eQTL-based)^31^, EndMT pathway membership (Hallmark EMT gene set)^26^, and key drivers of sex-specific endothelial CAD pathways^32^ were included as supportive evidence and assigned lower weights (0.25). Weights were summed across evidence categories to generate a cumulative prioritization score, highlighting genes supported by multiple orthogonal lines of evidence across human transcriptomics, genetics, experimental models, and network analysis.

## Results

### EndMT is Induced in Both Male and Female Endothelial Cells

Endothelial cells (ECs) isolated from the ascending aortas of healthy five male and five female heart transplant donors, along with ECs isolated from the atherosclerotic lesions of three male and six female patients undergoing carotid endarterectomy were subjected to the EndMT assay using TNFα, TGFβ, or a combined TNFα+TGFβ treatment (Figure 1). All conditions produced characteristic morphological changes, with cells shifting from a cobblestone endothelial appearance to an elongated, spindle-like mesenchymal morphology in both male and female ECs. Immunofluorescence staining showed a decrease in the endothelial marker PECAM1 and an increase in the mesenchymal marker FN1 across both sexes (Figure 1B).

Bulk RNA-seq was performed for all donors across all stimulation conditions and time points (0, 24, 72, and 144 hours). Principal component analysis (PCA) revealed clear separation by treatment stimulation primarily for TNFα (Supplemental Figure S2A-B). Sex contributed modestly to the global transcriptomic variance, with partial separation along PC3 and PC4, particularly in the healthy ECs (Supplemental Figure S2C-D). Expression of canonical endothelial and mesenchymal markers changed dynamically over the course of stimulation. TNFα stimulation was supported by elevated *NFKB1* expression, and TGFβ activity was confirmed by increased *SNAI1* levels (Supplemental Figure 1B-C). *PECAM1* expression declined in both male and female ECs irrespective of disease status (Figure 1C & E), with TGFβ alone producing a stronger reduction in atherosclerotic ECs. Additional endothelial markers such as *CDH5, NOS3*, and *VWF* showed similar time-dependent decreases (Supplemental Figure S1D-F). Conversely, the mesenchymal marker *FN1* increased over time in both healthy and atherosclerotic ECs from males and females (Figure 1D & F). Other mesenchymal markers including *ACTA2, CDH2*, and *TAGLN*, followed the same pattern (Supplemental Figure S1G-I).

### Sex-Specific EndMT Regulation Differs Between Healthy and Atherosclerotic Endothelial Cells

To test our hypothesis that EndMT is regulated in a sex-specific manner, we conducted a sex-specific trajectory analysis of differences in transcriptional regulation between males and females within each treatment condition. We focused on whether males and females exhibited distinct EndMT trajectories over time (Figure 2A), including: (i) directional differences, where the sexes show opposite temporal expression patterns; (ii) sex-biased effects, where only one sex shows up- or downregulation; and (iii) magnitude differences, where both sexes respond but to different degrees. Sex-specific genes were defined using an FDR < 0.25, with positive logFC indicating higher expression in males and negative values indicating higher expression in females. In healthy ECs, the sex-specific trajectory analysis identified 3,492 unique sex-biased genes across all conditions: 12 in control, 507 in TNFα, 4 in TGFβ, and 3,268 in the combined TGFβ+TNFα treatment. A number of these genes overlapped between conditions (Figure 2B). This indicated a strong sex-specific regulation in healthy ECs. In contrast, atherosclerotic ECs showed far fewer sex differences, with only 30 unique sex-biased genes identified: 7 in control, 7 in TNFα, 3 in TGFβ, and 22 in TGFβ+TNFα (Figure 2C).

**Figure 2.**
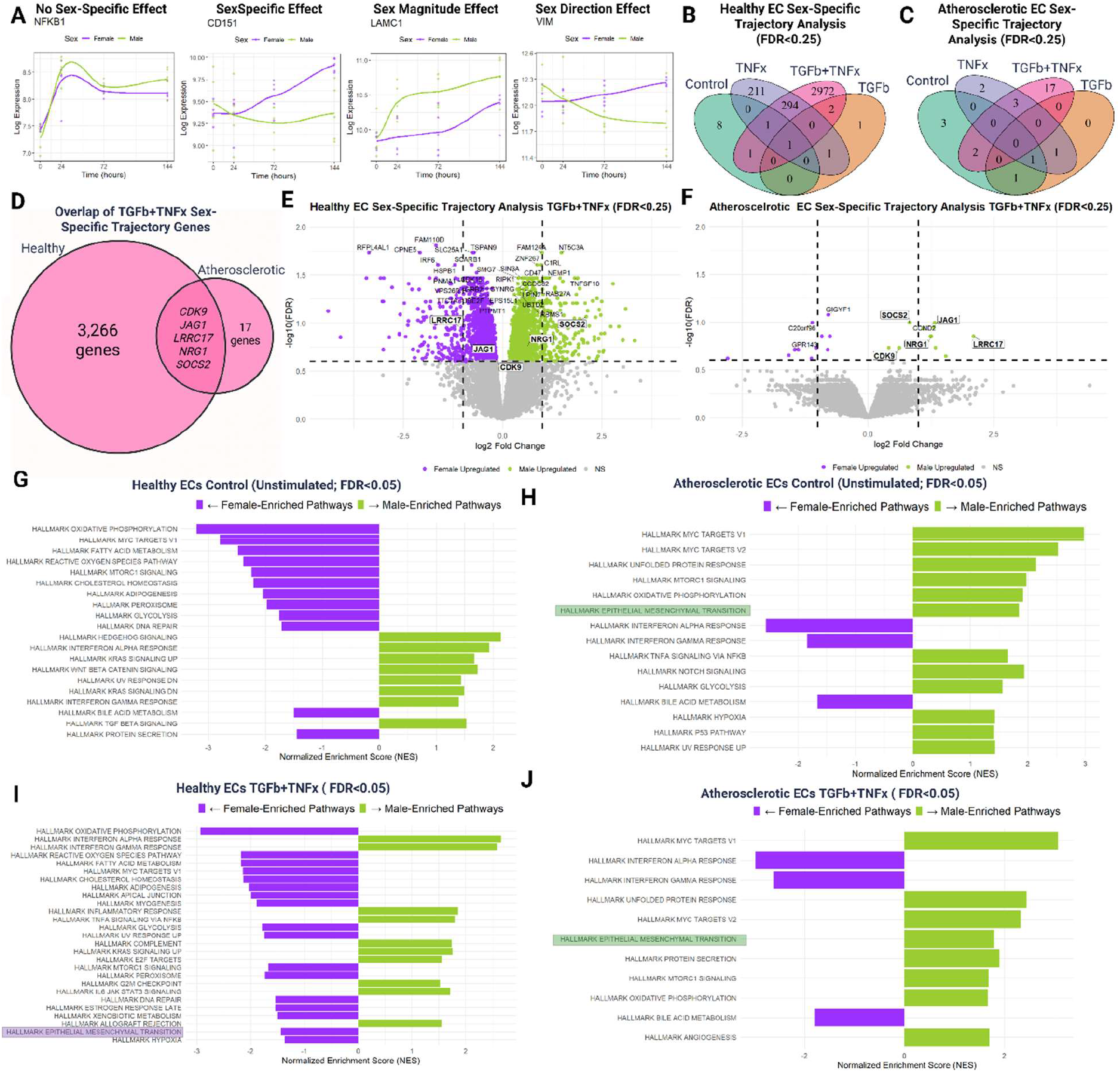
RNA-seq analysis of the in vitro EndMT assay. (A) Examples of representative sex-specific trajectories from the healthy EC TGFβ+TNFα condition. (B) Sex-specific trajectory analysis for each treatment (Control, TGFβ, TNFα, and TGFβ+TNFα), identifying 3,492 sex-specific genes in healthy ECs (FDR < 0.25). (C) Sex-specific trajectory analysis for atherosclerotic ECs, which show markedly reduced sex-specific regulation, with only 30 sex-specific genes detected (FDR < 0.25). (D) The TGFβ+TNFα condition produced the strongest sex-specific gene regulation in both vascular beds. Comparing the 3,271 sex-biased genes from healthy ECs with the 22 sex-biased genes from atherosclerotic ECs revealed five shared genes: *CDK9, JAG1, LRRC17, NRG1*, and *SOCS2*. (E–F) Volcano plots of sex-specific genes from the TGFβ+TNFα condition for healthy (E) and atherosclerotic (F) ECs. (G–J) Pathway enrichment analysis for Control and TGFβ+TNFα conditions for healthy and atherosclerotic ECs (FDR < 0.05).

In both healthy and atherosclerotic ECs, the combined TGFβ+TNFα stimulation produced the largest sex-specific differences, possibly by activation of EndMT through two distinct pathways. Also, TNFα alone showed a stronger sex effect than TGFβ alone. Comparing the 3,268 sex biased genes from the healthy EC TGFβ+TNFα condition with the 22 sex-biased genes from the atherosclerotic ECs revealed only 5 overlapping genes (Figure 2C). Volcano plots for the TGFβ+TNFα condition further illustrate the pronounced sex-specific responses in healthy ECs compared to atherosclerotic ECs (Figure 2D–E). Complete gene-level results for the sex-specific trajectory analysis models are available in the Supplemental Tables 2-9.

We also performed pathway enrichment analysis for the sex-specific trajectory analysis models in the control and TGFβ+TNFα conditions using an unbiased ranked approach based on the t-statistic. In healthy control ECs, we found numerous sex-divergent pathways, indicating baseline sex differences in endothelial regulation (Figure 2F). Under our contrast specification (male minus female), positive logFC and t-statistics indicate higher expression in males, whereas negative values indicate higher expression in females. Pathways were considered significantly enriched at FDR < 0.05. EndMT-related activation was not detected in the control condition because these cells were not stimulated with TGFβ+TNFα. Although there is no formal gene ontology term for EndMT, the Hallmark epithelial-to-mesenchymal transition (EMT) pathway is commonly used as a proxy in the literature, as EMT regulation in cancer shares many molecular features with EndMT^8^. However, in atherosclerotic ECs, EndMT-related pathways were already higher in males even under control conditions, indicating that they were more vulnerable for EndMT activation without any biochemical stimulus (Figure 2G). In contrast, under TGFβ+TNFα stimulation, healthy females showed greater EndMT pathway activation (Figure 2H). In atherosclerotic ECs, males continued to exhibit the stronger response (Figure 2I). This sex difference in pathway dominance between stimulated healthy and atherosclerotic ECs was not limited to EndMT-related programs but was also extended to Myc Targets, interferon-α, interferon-γ, and oxidative phosphorylation pathways. These findings suggest that sex-specific regulation of EndMT may change over time and with disease progression.

### Human EC Sex-Specific Pseudotime Trajectories Support Sex-Specific Patterns of EndMT Activation

To determine whether the sex-specific transcriptional programs observed in our in vitro EndMT assay are reflected in a pathophysiological context, we performed a sex-specific pseudotime trajectory analysis using three publicly available human carotid plaque datasets from 36 male and 28 female patients^6,27,28^. Integration of three independent datasets yielded 1,815 ECs (Figure 3A), including 918 female and 897 male cells (Figure 3B). Clustering was used to identify the starting point for trajectory analysis (Figure 3C). Based on canonical endothelial, mesenchymal, inflammatory, and smooth muscle markers, EC1 was designated as the root state (Figure 3D). This marker-based evaluation also supported the removal of EC7 as an inflammatory population and EC12 as an SMC- or fibroblast-like population.

**Figure 3:**
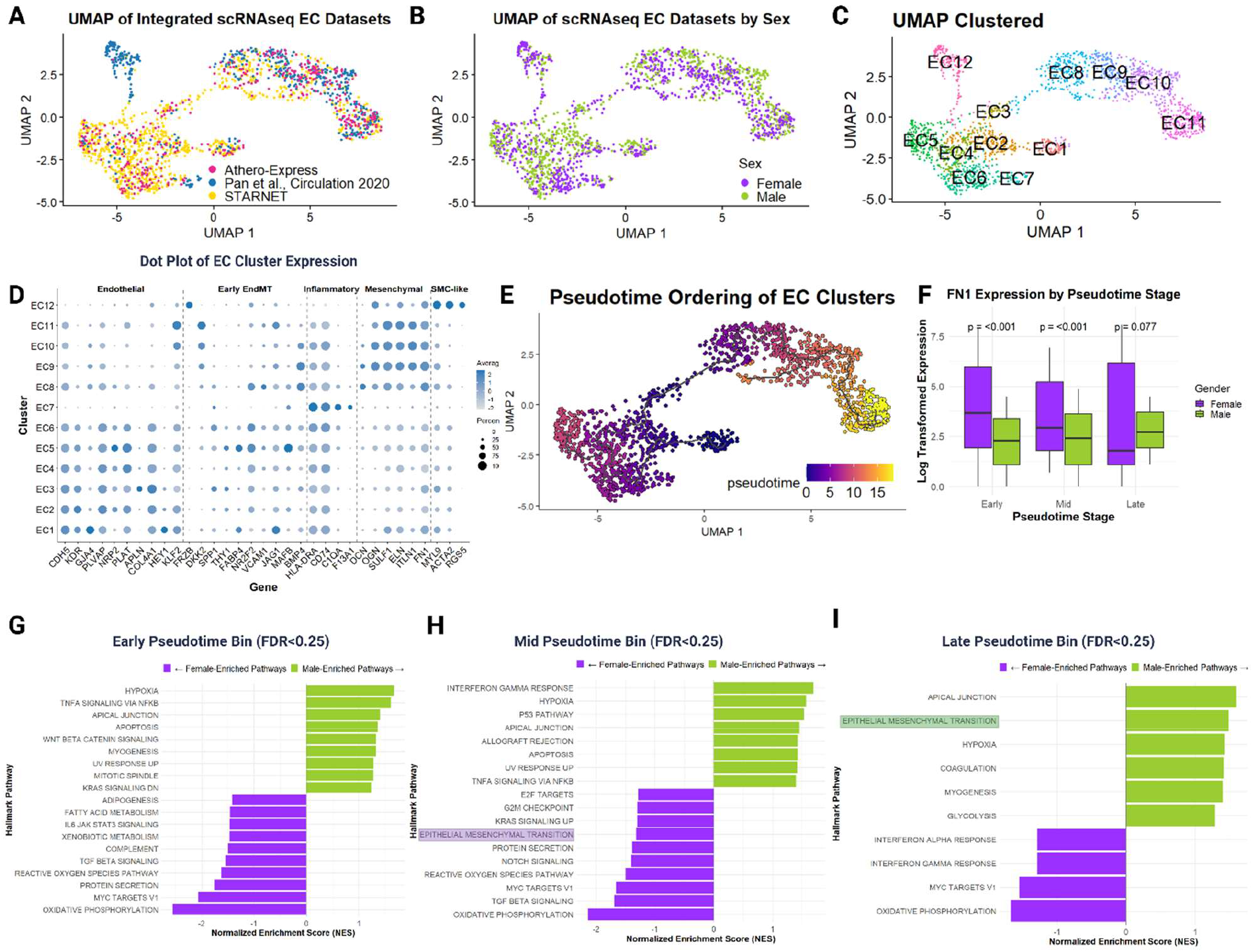
scRNA-seq sex-specific pseudotime trajectory analysis of human ECs. Three independent scRNA-seq datasets were integrated for pseudotime reconstruction. (A) UMAP showing the distribution of cells from each dataset. (B) UMAP colored by sex. (C) UMAP with cluster identities. (D) Dot plot displaying average expression of endothelial and mesenchymal marker genes across clusters, used to define Cluster 1 as the pseudotime root and to exclude Clusters 7 and 12 as non-EC populations. (E) Pseudotime trajectory progression through clusters. (F) *FN1* expression differences between males and females for early, mid, and late pseudotime bins. (G–I) Pathway enrichment analyses for early, mid, and late pseudotime bins (FDR < 0.25).

Pseudotime analysis revealed a continuous endothelial-to-mesenchymal progression (Figure 3E). Graph-based differential expression testing identified 2,206 pseudotime associated genes, 1,107 sex-associated genes, and 257 genes influenced by the interaction between pseudotime and sex (Supplemental Tables 10-12). Notably, sex accounted for a substantial proportion of transcriptional variance along the trajectory, indicating that male and female ECs take partially distinct transcriptional routes through the EndMT continuum. Full differential expression results are reported in the Supplemental Tables 10-12

We next compared the pseudotime-by-sex gene set with our in vitro sex-specific trajectory findings. Forty-one genes overlapped with the healthy EC TGFβ+TNFα condition, whereas only one gene overlapped with the atherosclerotic EC TGFβ+TNFα condition. Although the overlap was modest, it suggests that endothelial cells exhibit greater transcriptional plasticity at pseudo-earlier stages of atherosclerotic progression compared to healthy aortic endothelial cells, which represent an earlier baseline state. This also suggests that the sex-specific programs observed in vitro capture, at least in part, fundamental features of early EndMT that are preserved in human carotid endothelium. This comparison also alleviated concerns that the strong differences observed between healthy and atherosclerotic ECs in vitro were simply due to vascular bed differences, as the carotid scRNA-seq data aligned more closely with the healthy aortic EC responses. While vascular bed–specific differences cannot be fully excluded, alignment with carotid EC scRNA-seq supports disease stage rather than vascular bed origin as the dominant driver.

Interestingly, mirroring our in vitro findings, females exhibited stronger EndMT activation at earlier pseudotime stages (similar to healthy ECs), whereas males showed greater activation at later stages (similar to atherosclerotic ECs). This pattern was evident in the expression trajectory of *FN1*, a canonical mesenchymal marker (Figure 3F), and was further supported by pathway enrichment analysis (Figure 3G-I; FDR < 0.25). These data indicate that sex-specific EndMT regulation observed under controlled perturbation conditions is also detectable in human carotid endothelium, reinforcing the relevance of these programs to in vivo plaque biology.

### WGCNA Identifies EndMT-Associated Gene Modules with Sex- and Disease-Dependent Preservation

Weighted gene co-expression network analysis (WGCNA) was performed using gene expression data from healthy female endothelial cells subjected to TGFβ + TNFα stimulation at 24, 72, and 144 hours. Network analysis identified multiple co-expression modules with distinct eigengene correlation structure, with the gene dendrogram and module assignments derived from the healthy female reference network shown in the Supplemental Figure 3A. To prioritize modules relevant to EndMT, modules were ranked using a composite EndMT module score that integrated the strength of module eigengene association with an EMT score and enrichment for Hallmark EMT genes. This analysis identified the magenta module as the top-ranked EndMT-associated module, followed by the dark green module, with both showing substantially higher EndMT scores than all remaining modules (Figure 4A).

**Figure 4.**
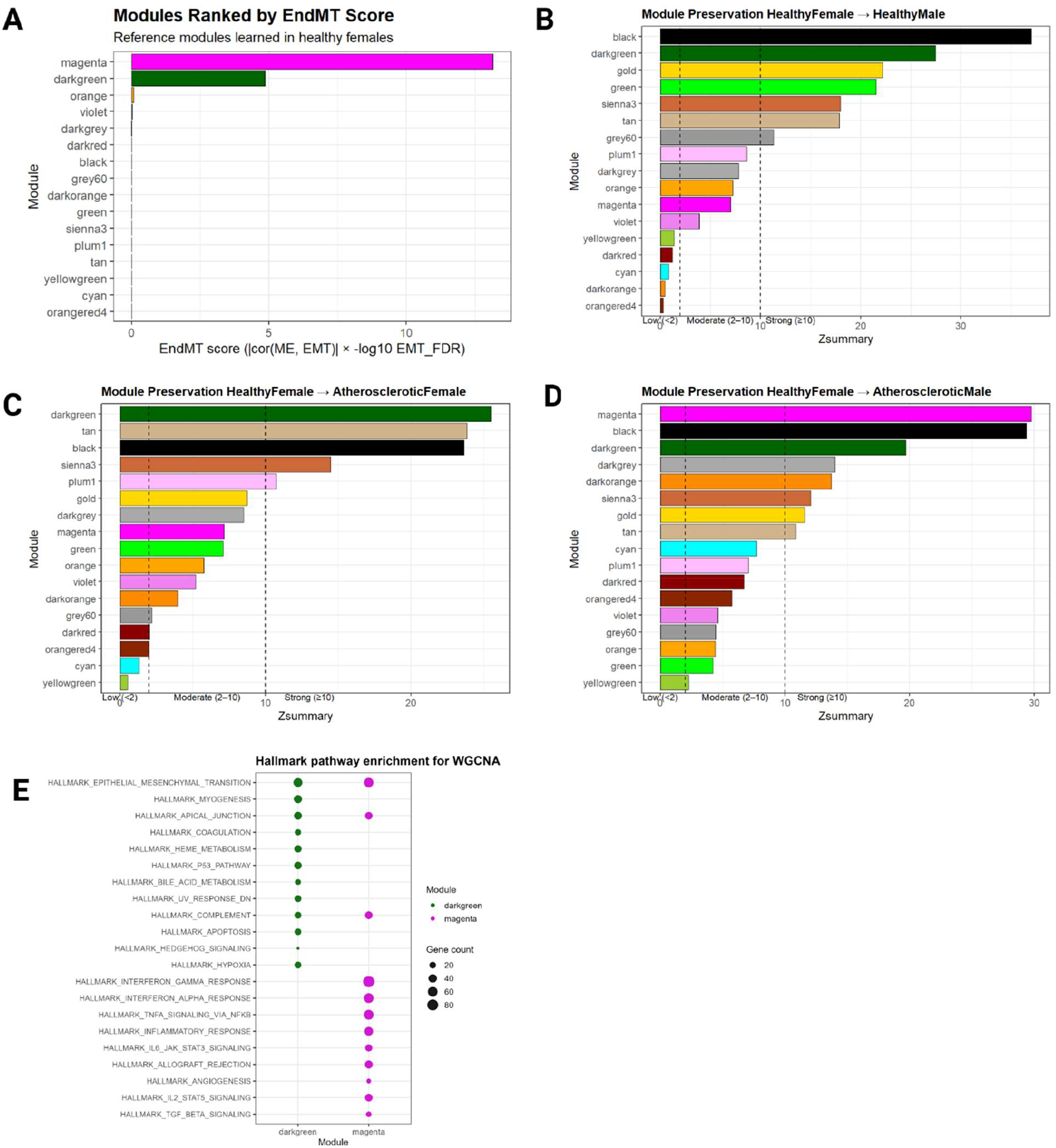
Preservation and characterization of EndMT-associated WGCNA modules. (A) Weighted gene co-expression network analysis (WGCNA) modules learned in healthy female endothelial cells ranked by an EndMT score. (B–D) Module preservation analysis of healthy female reference modules assessed in (B) healthy male, (C) atherosclerotic female, and (D) atherosclerotic male endothelial cells. Dashed lines denote standard WGCNA preservation thresholds: Zsummary < 2 (low preservation), 2–10 (moderate preservation), and ≥10 (strong preservation). (E) Over-representation analysis of Hallmark gene sets for the magenta and dark green modules. Dot size indicates the number of overlapping genes between each module and the pathway.

To assess the robustness of these EndMT-associated modules across sex and disease contexts, module preservation was evaluated using the healthy female network as the reference. When tested in healthy male endothelial cells, several modules exhibited strong preservation, including the dark green module, whereas the magenta module showed reduced preservation relative to other highly ranked modules (Figure 4B). In atherosclerotic females, the dark green module remained strongly preserved, while the magenta module exhibited only moderate preservation (Figure 4C). Notably, in atherosclerotic males, both the magenta and dark green modules showed strong preservation with the healthy female reference (Figure 4D). These results indicate that preservation of EndMT-associated transcriptional programs varies by both sex and disease state. The dark green module was consistently preserved across healthy and atherosclerotic contexts in both sexes, suggesting it represents a stable EndMT-associated transcriptional program. All module EndMT score and preservation results can be found in Supplemental Table 13. In contrast, preservation of the magenta module was more context dependent, varying by sex and disease state, consistent with a more dynamic or condition-sensitive EndMT program.

To further assess the robustness of sex-specific differences in module preservation and to account for unequal sample sizes between atherosclerotic females (n = 6) and males (n = 3), a balanced resampling analysis was performed using equal numbers of atherosclerotic male and female samples, with preservation recalculated across 50 iterations (20 permutations per iteration). Consistent with the single-run preservation analysis, the dark green module showed stable preservation across resampling iterations in both sexes, whereas preservation of the magenta module was highly sex dependent, with consistently low preservation in atherosclerotic females and strong preservation in atherosclerotic males (Supplemental Figure 3B). These findings indicate that sex differences in magenta module preservation are not driven by sample size imbalance but reflect a reproducible, sex-specific difference in network preservation. The genes in the magenta and darkgreen modules can be found in Supplemental Table 14.

Pathway enrichment analysis revealed limited overlap between EndMT-associated modules, indicating that they capture distinct components of the EndMT transcriptional response (Supplemental Table 15). The three pathways shared between the magenta and dark green modules: EMT, apical junction, and complement represent core features of endothelial plasticity. In contrast, the remaining enriched pathways were module specific. The magenta module was enriched for inflammatory, cytokine, angiogenic, and TGFβ-associated pathways, consistent with a more dynamic EndMT program, whereas the dark green module was enriched for pathways related to coagulation, hypoxia, p53 signaling, and metabolic homeostasis, consistent with a more stable, baseline EndMT-associated transcriptional program.

### Prioritization of Top Sex-specific Regulators of EndMT

To identify high confidence sex-specific regulators of EndMT, we implemented a weighted gene-prioritization framework to rank genes identified in our sex-specific trajectory analyses. In the human datasets, 3,275 genes exhibited sex-specific trajectory differences in healthy endothelial cell (EC) RNA-seq under TGFβ+TNFα conditions, whereas 22 genes showed sex-specific differences in atherosclerotic EC RNA-seq (FDR < 0.25; Supplemental Tables 5 and 9). In addition, 183 genes were identified through sex-specific pseudotime analysis of human single-cell RNA-seq data (FDR < 0.25; Supplemental Table 12). In total, 3,990 unique genes were identified across these sex-specific EndMT trajectory datasets. There were no genes identified across all three independent experimental datasets, which is not unexpected given their distinct biological origins and experimental contexts.

To further validate our findings and increase confidence in the prioritized candidates, we integrated evidence from published genetic studies of coronary artery disease (CAD). Specifically, we assessed overlap with genes implicated in CAD risk by large-scale genome-wide association studies (GWAS). This included the 897 CAD-associated genes reported by Aragam et al., derived from analyses of more than one million participants and representing a comprehensive genetic framework for CAD susceptibility. In addition, we incorporated 258 CAD GWAS genes identified by Tcheandjieu et al., which leveraged multi-ancestry association mapping to refine candidate causal genes^30^. Of these, 153 genes overlapped with the genes identified in our sex-specific EndMT trajectory analyses. Additionally, we incorporated the 11 genes reported in that study to exhibit sex-biased genetic effects on CAD risk; among these, only *COL4A1* was identified in our dataset^29^. We further incorporated endothelial-specific systems genetics evidence by assessing overlap with 92 genes identified through expression quantitative trait locus (eQTL) mapping in primary human aortic endothelial cells from Stolze et al.^31^; in total, 23 of these genes overlapped with our dataset. In addition, we assessed overlap with the MSigDB Hallmark epithelial mesenchymal transition (EMT) gene set, which comprises 200 curated genes, of which 122 overlapped with our sex-specific EndMT trajectory dataset^26^.

To incorporate network-level regulatory evidence, we leveraged key-driver genes identified in sex-stratified gene regulatory networks from Hartman et al., who constructed causal networks from human atherosclerotic plaque transcriptomes to identify sex-specific drivers of vascular disease. In that study, key driver analysis identified 51 genes within a female-biased, endothelial-enriched yellow module associated with vascular remodeling and disease progression^32^. Of these, 11 key driver genes overlapped with the genes identified in our sex-specific EndMT trajectory analyses, providing independent network-level support for a subset of prioritized candidates. Finally, as an additional line of evidence, we incorporated network-level support from our WGCNA, integrating genes from the sex-biased EndMT-associated magenta module (Supplemental Table 13) to further refine and prioritize sex-specific EndMT regulators, 336 of these genes were present in our data set.

Each of the 3,990 genes was assigned a cumulative weighted score based on support across sex-specific trajectory datasets, genetic association studies, and network-level analyses, and genes were ranked according to this integrated score (Supplemental Table 17). The top 10 prioritized genes are summarized in Table 1 and include *COL4A1, PECAM1, CD151, JAG1, FN1, NEDD9, PODXL, MAFB, PROCR*, and *CDH13*. Among the highest-ranked genes, several represent putative upstream regulators of sex-specific EndMT programs, while others (e.g., *PECAM1, FN1*) correspond to established endothelial or mesenchymal state markers, providing internal biological validation of the prioritization framework.

## Discussion

In this study, we integrated bulk and single-cell transcriptomic analyses from human endothelial cells to define sex-specific differences in EndMT regulation during atherosclerosis. The role of EndMT in atherosclerotic disease remains controversial^8,13,33^. Early studies largely characterized EndMT as a pathogenic process, associating mesenchymal marker expression with fibrosis, inflammation, and plaque progression^34^. More recent literature, however, suggests that EndMT may also serve adaptive or protective functions, particularly during early stages of disease^8^. We hypothesized that EndMT regulation exhibits sex-specific differences that may be associated with known clinical sex differences in plaque biology, including variation in plaque stability, fibrous cap integrity, and susceptibility to erosion versus rupture.

Our findings support a stage dependent and sex-specific model of EndMT regulation, in which endothelial cells engage adaptive or maladaptive mechanisms depending on the timing and extent of activation. EndMT pathway activation was more prominent in females during early disease stages, a pattern consistently observed across both in vitro EndMT assays and scRNA-seq pseudotime analyses. These sex-biased early EndMT programs align with clinical observations that younger females exhibit lower rates of plaque rupture, although plaque erosion remains a relevant disease manifestation^2^. Prior studies have linked *FN1* associated EndMT states to endothelial activation during early plaque development, consistent with mechanisms implicated in plaque erosion, which is characterized by endothelial dysfunction, inflammation, and detachment rather than fibrous cap rupture^13,35^. Consistent with this literature, we observed higher *FN1* expression in females during early and intermediate disease stages. However, this sex difference diminished as disease progressed, with increased EndMT signatures observed in males at later stages, reflecting the greater plaque instability observed in both sexes with advancing age^36,37^. Although we did not perform lineage tracing of endothelial cells in this study, prior lineage-tracing analyses in mouse models have similarly reported increased EndMT in males during late-stage disease^4^. Consistent with this stage- and sex-dependent model, network-level analysis using WGCNA revealed that EndMT regulation is organized into distinct transcriptional programs with differing sensitivity to sex and disease context. The top EndMT-associated module identified in healthy endothelial cells was enriched for inflammatory and mesenchymal signaling pathways and showed marked sex-dependent differences in temporal activation. These results indicate that sex-specific regulation of EndMT varies across disease stage, supporting a model in which EndMT may contribute to adaptive remodeling early in disease but becomes maladaptive when excessively activated or improperly regulated.

Similar attenuation of sex differences with disease progression has been observed in many other disease contexts, often involving shared cellular stress-response and phenotypic plasticity programs analogous to EndMT. In heart failure, sex-biased differences in disease prevalence, cardiac remodeling, and inflammatory signaling are most evident early in disease and tend to converge as pathology advances, coinciding with activation of common fibrotic and endothelial stress programs^38–40^. Comparable patterns have been described in autoimmune and inflammatory disorders, where sex-biased susceptibility and immune activation are strongest early in disease and diminish as chronic pathology becomes established^41^. Likewise, in metabolic diseases such as insulin resistance and type 2 diabetes, sex differences in vascular and inflammatory responses are evident during early dysfunction but converge with increasing disease severity^42^. Together, these observations underscore the importance of disease stage when interpreting sex-specific mechanisms and support the concept that sex differences in EndMT and related endothelial plasticity programs are most pronounced early in disease, helping to explain why clinical sex differences often attenuate with disease progression.

Our gene-prioritization framework identified a set of top sex-specific EndMT regulators, highlighting multiple entry points through which biological sex may shape EndMT trajectories. Several candidates including *COL4A1, PECAM1, CD151, PODXL, FN1* and *CDH13* are closely linked to endothelial junctional integrity, basement membrane organization, extracellular matrix interactions, and shear-stress sensing, supporting a model in which sex differences in biomechanical signaling influence early EndMT activation^43–46^. Notably, *COL4A1* has also been identified as a sex-biased locus in cardiovascular genome-wide association studies, providing strong independent human genetic support for its role in sex-specific vascular disease mechanisms^29^. While we did not observe a significant sex difference in *COL4A1* expression in our biochemically induced in vitro EndMT assays, this sex-specific effect was evident in vivo, suggesting that *COL4A1*-mediated sex differences may be driven by biomechanical cues not fully recapitulated in static culture systems. Consistent with this interpretation, a recent preprint using flow-induced EndMT reported sex-dependent regulation of *COL4A1*, further supporting a role for hemodynamic context in uncovering sex-specific EndMT regulation^5^.

Beyond regulators linked to extracellular matrix organization and biomechanical sensing, our prioritized gene set also highlights sex-specific differences in intracellular signaling, cytoskeletal remodeling, and stress-responsive pathways that shape EndMT trajectories. *JAG1*, a key Notch ligand, implicates cell-cell communication and fate signaling in sex-specific EndMT regulation, potentially modulating the balance between endothelial maintenance and mesenchymal activation^47^. In parallel, regulators associated with cytoskeletal remodeling and migratory signaling (*NEDD9, FN1*, and *CDH13*) point to sex differences in endothelial plasticity and inflammatory activation states, consistent with partial EndMT programs^48–50^. Concurrent modulation of canonical endothelial markers such as *PECAM1* further reinforces the presence of intermediate EndMT states rather than complete loss of endothelial identity^51^. Additional candidates, including *PROCR* and *MAFB*, link sex-specific EndMT regulation to anticoagulant signaling and macrophage-associated transcriptional programs, suggesting that immune-vascular crosstalk may intersect with endothelial plasticity in a sex-dependent manner^29,52^. Collectively, these findings indicate that sex-specific EndMT regulation does not arise from a single dominant pathway but instead reflects coordinated differences across mechanotransduction, junctional signaling, cytoskeletal dynamics, and inflammatory responsiveness that may bias vascular remodeling toward adaptive or maladaptive plaque phenotypes.

Several limitations should be acknowledged. There is currently no standardized method to induce EndMT in vitro, and commonly used biochemical models may fail to capture key physiological drivers such as disturbed flow, hypoxia, and mechanical strain^53^. In addition, pseudotime analyses are inherently model-dependent and rely on assumptions about cellular progression that may not fully reflect in vivo dynamics ^54^. Finally, while our integrative approach identifies high-confidence sex-specific regulators of EndMT, functional validation is required to establish causal roles and directly link these programs to plaque stability. Future studies will focus on functional validation of top sex-specific EndMT regulators using targeted perturbation approaches in primary human endothelial cells to test whether sex-biased EndMT programs differentially influence endothelial functions relevant to plaque stability. Complementary in vivo lineage-tracing and conditional knockout models combined with snRNA-seq will be used to determine whether sex-specific EndMT trajectories causally contribute to differences in plaque composition and stability. These approaches will be critical for distinguishing adaptive versus maladaptive EndMT trajectories and for identifying therapeutically actionable windows and molecular targets through which modulation of EndMT plasticity may be most effective

## References

1. Covani, M. et al. Plaque Vulnerability and Cardiovascular Risk Factor Burden in Acute Coronary Syndrome: An Optical Coherence Tomography Analysis. J. Am. Coll. Cardiol. 86, 77–89 (2025).

2. Seegers, L. M. et al. Sex Differences in Culprit Plaque Characteristics Among Different Age Groups in Patients With Acute Coronary Syndromes. Circ. Cardiovasc. Interv. 15, e011612 (2022).

3. Sakkers, T. R. et al. Sex differences in the genetic and molecular mechanisms of coronary artery disease. Atherosclerosis 384, 117279 (2023).

4. Shin, J. et al. Unraveling the Role of Sex in Endothelial Cell Dysfunction: Evidence From Lineage Tracing Mice and Cultured Cells. Arterioscler. Thromb. Vasc. Biol. 44, 238–253 (2024).

5. Tardajos-Ayllon, B. et al. TWIST1 drives endothelial-to-mesenchymal-transition to stabilize atherosclerotic plaques. 2025.05.19.654847 Preprint at 10.1101/2025.05.19.654847 (2025).

6. Sukhavasi, K. et al. Single-cell RNA sequencing reveals sex differences in the subcellular composition and associated gene-regulatory network activity of human carotid plaques. Nat. Cardiovasc. Res. 4, 412–432 (2025).

7. Slenders, L. et al. Endothelial-to-mesenchymal transition gene signature derived from single-cell transcriptomics of human atherosclerotic tissue associates with stable plaque histological characteristics. Vascul. Pharmacol. 159, 107498 (2025).

8. Hall, I. F., Aikawa, E., Sluimer, J., Baker, A. H. & Kovacic, J. C. Endothelial to mesenchymal transition in cardiovascular diseases: molecular insights and clinical perspectives. Eur. Heart J. ehaf670 (2025) doi:10.1093/eurheartj/ehaf670.

9. Kovacic, J. C. et al. Endothelial to Mesenchymal Transition in Cardiovascular Disease: JACC State-of-the-Art Review. J. Am. Coll. Cardiol. 73, 190–209 (2019).

10. Evrard, S. M. et al. Endothelial to mesenchymal transition is common in atherosclerotic lesions and is associated with plaque instability. Nat. Commun. 7, 11853 (2016).

11. Chen, P.-Y. et al. Endothelial TGF-β signalling drives vascular inflammation and atherosclerosis. Nat. Metab. 1, 912–926 (2019).

12. Tombor, L. S. et al. Single cell sequencing reveals endothelial plasticity with transient mesenchymal activation after myocardial infarction. Nat. Commun. 12, 681 (2021).

13. Gole, S., Tkachenko, S., Masannat, T., Baylis, R. A. & Cherepanova, O. A. Endothelial-to-Mesenchymal Transition in Atherosclerosis: Friend or Foe? Cells 11, 2946 (2022).

14. Newman, A. A. et al. Multiple cell types contribute to the atherosclerotic lesion fibrous cap by PDGFRβ and bioenergetic mechanisms. Nat. Metab. 3, 166–181 (2021).

15. Hall, I. F., Kishta, F., Xu, Y., Baker, A. H. & Kovacic, J. C. Endothelial to mesenchymal transition: at the axis of cardiovascular health and disease. Cardiovasc. Res. 120, 223–236 (2024).

16. Qian, C. et al. Broadening horizons: molecular mechanisms and disease implications of endothelial-to-mesenchymal transition. Cell Commun. Signal. 23, 16 (2025).

17. Yoshimatsu, Y. & Watabe, T. Emerging roles of inflammation-mediated endothelial–mesenchymal transition in health and disease. Inflamm. Regen. 42, 9 (2022).

18. Kempe, S., Kestler, H., Lasar, A. & Wirth, T. NF-κB controls the global pro-inflammatory response in endothelial cells: evidence for the regulation of a pro-atherogenic program. Nucleic Acids Res. 33, 5308–5319 (2005).

19. Robert, J. Sex differences in vascular endothelial cells. Atherosclerosis 384, 117278 (2023).

20. Lu, C. et al. Sex-specific differences in cytokine signaling pathways in circulating monocytes of cardiovascular disease patients. Atherosclerosis 384, (2023).

21. Alvandi, Z. & Bischoff, J. Endothelial-Mesenchymal Transition in Cardiovascular Disease. Arterioscler. Thromb. Vasc. Biol. 41, 2357–2369 (2021).

22. Navab, M. et al. Monocyte migration into the subendothelial space of a coculture of adult human aortic endothelial and smooth muscle cells. J. Clin. Invest. 82, 1853–1863 (1988).

23. Jawitz, O. K. et al. Donor and recipient age matching in heart transplantation: Analysis of the UNOS Registry. Transpl. Int. Off. J. Eur. Soc. Organ Transplant. 32, 1194–1202 (2019).

24. Sakkers, T. R. et al. Atherosclerotic fibrous plaques in females are characterized by endothelial-to-mesenchymal transition and linked to smoking. 2024.10.01.24314739 Preprint at 10.1101/2024.10.01.24314739 (2024).

25. Diez Benavente, E. et al. Atherosclerotic Plaque Epigenetic Age Acceleration Predicts a Poor Prognosis and Is Associated With Endothelial-to-Mesenchymal Transition in Humans. Arterioscler. Thromb. Vasc. Biol. 44, 1419–1431 (2024).

26. HALLMARK_EPITHELIAL_MESENCHYMAL_TRANSITION. https://www.gsea-msigdb.org/gsea/msigdb/cards/HALLMARK_EPITHELIAL_MESENCHYMAL_TRANSITION.html.

27. Diez Benavente, E. et al. Female Gene Networks Are Expressed in Myofibroblast-Like Smooth Muscle Cells in Vulnerable Atherosclerotic Plaques. Arterioscler. Thromb. Vasc. Biol. 43, 1836–1850 (2023).

28. Pan, H. et al. Single-Cell Genomics Reveals a Novel Cell State During Smooth Muscle Cell Phenotypic Switching and Potential Therapeutic Targets for Atherosclerosis in Mouse and Human. Circulation 142, 2060–2075 (2020).

29. Aragam, K. G. et al. Discovery and systematic characterization of risk variants and genes for coronary artery disease in over a million participants. Nat. Genet. 54, 1803–1815 (2022).

30. Tcheandjieu, C. et al. Large-scale genome-wide association study of coronary artery disease in genetically diverse populations. Nat. Med. 28, 1679–1692 (2022).

31. Stolze, L. K. et al. Systems Genetics in Human Endothelial Cells Identifies Non-coding Variants Modifying Enhancers, Expression, and Complex Disease Traits. Am. J. Hum. Genet. 106, 748–763 (2020).

32. Hartman, R. J. G. et al. Sex-Stratified Gene Regulatory Networks Reveal Female Key Driver Genes of Atherosclerosis Involved in Smooth Muscle Cell Phenotype Switching. Circulation 143, 713–726 (2021).

33. Simons, M. Endothelial-to-mesenchymal transition: advances and controversies. Curr. Opin. Physiol. 34, 100678 (2023).

34. Kovacic, J. C., Mercader, N., Torres, M., Boehm, M. & Fuster, V. Epithelial-to-Mesenchymal and Endothelial-to-Mesenchymal Transition. Circulation 125, 1795–1808 (2012).

35. Al-Yafeai, Z. et al. Endothelial FN (Fibronectin) Deposition by α5β1 Integrins Drives Atherogenic Inflammation. Arterioscler. Thromb. Vasc. Biol. 38, 2601–2614 (2018).

36. Wang, J. C. & Bennett, M. Aging and Atherosclerosis. Circ. Res. 111, 245–259 (2012).

37. Bakker, M. de et al. The age- and sex-specific composition of atherosclerotic plaques in vascular surgery patients. Atherosclerosis 310, 1–10 (2020).

38. Martin, T. G. & Leinwand, L. A. Hearts apart: sex differences in cardiac remodeling in health and disease. J. Clin. Invest. 134, e180074.

39. Watts, K. M., Nichols, W. & Richardson, W. J. Computational screen for sex-specific drug effects in a cardiac fibroblast signaling network model. Sci. Rep. 13, 17068 (2023).

40. Watts, K. & Richardson, W. J. Effects of Sex and 17 β-Estradiol on Cardiac Fibroblast Morphology and Signaling Activities In Vitro. Cells 10, 2564 (2021).

41. Fairweather, D., Frisancho-Kiss, S. & Rose, N. R. Sex Differences in Autoimmune Disease from a Pathological Perspective. Am. J. Pathol. 173, 600–609 (2008).

42. Kautzky-Willer, A., Leutner, M. & Harreiter, J. Sex differences in type 2 diabetes. Diabetologia 66, 986–1002 (2023).

43. Fleming, I., Fisslthaler, B., Dixit, M. & Busse, R. Role of PECAM-1 in the shear-stress-induced activation of Akt and the endothelial nitric oxide synthase (eNOS) in endothelial cells. J. Cell Sci. 118, 4103–4111 (2005).

44. Volonghi, I. et al. Role of COL4A1 in basement-membrane integrity and cerebral small-vessel disease. The COL4A1 stroke syndrome. Curr. Med. Chem. 17, 1317–1324 (2010).

45. Zhang, F. et al. Tetraspanin CD151 maintains vascular stability by balancing the forces of cell adhesion and cytoskeletal tension. Blood 118, 4274–4284 (2011).

46. Porras, G., Ayuso, M. S. & González-Manchón, C. Leukocyte-endothelial cell interaction is enhanced in podocalyxin-deficient mice. Int. J. Biochem. Cell Biol. 99, 72–79 (2018).

47. Souilhol, C. et al. JAG1-NOTCH4 mechanosensing drives atherosclerosis. Sci. Adv. 8, eabo7958.

48. Alba, G. A. et al. NEDD9 Is a Novel and Modifiable Mediator of Platelet-Endothelial Adhesion in the Pulmonary Circulation. Am. J. Respir. Crit. Care Med. 203, 1533–1545 (2021).

49. Ciszewski, W. M., Wawro, M. E., Sacewicz-Hofman, I. & Sobierajska, K. Cytoskeleton Reorganization in EndMT—The Role in Cancer and Fibrotic Diseases. Int. J. Mol. Sci. 22, 11607 (2021).

50. Rubina, K. et al. T-Cadherin (CDH13) and Non-Coding RNAs: The Crosstalk Between Health and Disease. Int. J. Mol. Sci. 26, 6127 (2025).

51. Lertkiatmongkol, P., Liao, D., Mei, H., Hu, Y. & Newman, P. J. Endothelial functions of PECAM-1 (CD31). Curr. Opin. Hematol. 23, 253–259 (2016).

52. Esmon, C. T. Inflammation and the activated protein C anticoagulant pathway. Semin. Thromb. Hemost. 32 Suppl 1, 49–60 (2006).

53. Islam, S. et al. The Mechanobiology of Endothelial-to-Mesenchymal Transition in Cardiovascular Disease. Front. Physiol. 12, (2021).

54. Hutton, A. & Meyer, J. G. Trajectory Inference for Single Cell Omics. ArXiv arXiv:2502.09354v1 (2025).

